# Pathogenesis of Alcohol-Exacerbated Malaria in *Plasmodium berghei*-Infected Mice

**DOI:** 10.64898/2026.04.30.720083

**Authors:** Bertrand Yuwong Wanyu, Emégam Nadège Kouémou, Methodius Shinyuy Lahngong, Helen Tiku Nda, Vanessa Tita Jugha, Fabrice Ambe Ngwa, Germain Sotoing Taiwe

## Abstract

**Introduction:** Malaria is still a pressing global health challenge, especially in sub-Saharan Africa, where behavioral factors such as alcohol consumption may exacerbate its impact. The present study is aimed at investigating the pathogenesis of alcohol-exacerbated malaria in *Plasmodium berghei*-infected an animal model (mice).

**Methods:** Male mice were separated into four treatment groups: control, alcohol control, *P. berghei* and *P. berghei* plus acute alcohol treatment groups. Animals were infected with malaria through intraperitoneal injection of *P. berghei* and an acute dose of ethanol (20% v/v) was introduced 48 hours post-infection. Parasitaemia was monitored using the Giemsa-stained thin blood smears. Haematological parameters were assessed using automated blood analyser. Liver function was evaluated by measuring serum levels of AST and ALT and cytokine profiles (TNF-α, INF-γ, IL-6, IL-1β) were quantified using ELISA kits.

**Results:** Results show that acute alcohol intake led to a significant increase in parasitaemia in the *P. berghei* group (p<0.01). Haematological analysis revealed a significant (p<0.001) reduction in RBC count, haemoglobin levels, haematocrit percentage, platelet count and others in the *P. berghei* plus acute alcohol group. Liver enzyme assays revealed an elevated AST and ALT levels (p<0.001) in the *P. berghei* group. Cytokine analysis revealed a significant (p<0.01) upregulation of pro-inflammatory cytokines (TNF-α INF-γ, IL-1β and IL-6), due to acute alcohol. These results suggest that alcohol exacerbates malaria pathogenesis by increasing parasitaemia, promoting immune dysregulation and liver injury, mediated by a shift toward a pro-inflammatory cytokine profile.

## 1. Introduction

Malaria, which is predominantly caused by the protozoan parasite *Plasmodium falciparum*, remains a public health burden, especially to sub-Saharan Africa, where it is responsible for a significant morbidity and mortality (1-3). The disease continues to inflict serious health challenges to the population, despite efforts to eradicate it. These challenges are exacerbated by complex interactions between the parasite, the host’s immune system and different socio-environmental factors. Alcohol is a frequently used and misused substance in sub-Saharan Africa (4, 5). Over 3 million deaths occur annually due to the harmful use of alcohol (6, 7). Over 5.1% of the global burden of disease is attributed to alcohol use, by World Health Organization (WHO) (8). The role of alcohol use in malaria pathogenesis is critical, and may influence disease progression significantly.

The pathogenesis of malaria involves a cascade of events, beginning with the parasite’s entry into the host’s bloodstream via the bite of an infected female *Anopheles* mosquito. From the bloodstream, the parasite moves to the liver, where in the hepatocytes, they undergo a pre-erythrocytic schizogonic cycle that leads to the production of merozoites (9, 10). The merozoites invades the red blood cells (RBCs) where they proliferate and cause a characteristic cyclical febrile episodes and haemolysis (11-13).

Both the innate and adaptive immunity are involved in the host immune response to *Plasmodium* infection (14, 15). Critical to this defence are cytokines such as TNF-α, IFN-γ, and IL-6, which help to control parasitaemia, but also contribute to tissue inflammation, particularly in severe forms of the disease (15-17). Acute alcohol consumption has been shown to modulate the immune system (18-20), potentially altering the host’s response to infectious diseases, such as malaria (21, 22). Ethanol, which is the primary ingredient in alcoholic beverages, interferes with the production of cytokines (23), the function of macrophages and T lymphocytes (24), and the integrity of the gut-liver axis, which plays a critical role in systemic immune regulation (25, 26). Acute ethanol exposure can suppress proinflammatory cytokines such as IFN-γ, which are essential for initiating and sustaining the Th1-type immune response necessary for managing liver and blood-stage *Plasmodium* infection by activating CD8+ T cells and promoting parasite elimination (27-29). This cytokine imbalance by alcohol can favour parasite survival and proliferation, which may lead to increase parasitaemia and exacerbated clinical symptoms (30, 31). In addition, alcohol intake has been shown to have a profound impact on liver functions, which is particularly relevant in malaria, as the liver plays a major role in the disease lifecycle. The primary mechanisms responsible for ethanol-induced liver damage are oxidative stress, mitochondrial dysfunction, and the activation of Kupffer cells, which are the liver’s resident macrophages (32, 33), which when activated release large amounts of reactive oxygen species (ROS) and pro-inflammatory cytokines (33, 34). As a result, blood levels of liver enzymes like AST and ALT are elevated, indicating hepatocellular damage (35, 36). Furthermore, ethanol intake aggravates hepatic dysfunction during malaria, where the parasite’s hepatic stage has already induced significant damages on the hepatocytes, resulting in a more severe liver damage (37).

Alcohol also aggravates malaria-induced haematological alterations. Ethanol consumption reduces the synthesis of RBCs by suppressing the bone marrow (38, 39), and worsens it by interfering with the integrity of the RBCs, making them more susceptible to risk of haemolysis during *Plasmodium* infection (40). The interaction between alcohol and malaria also extends to the gut-liver axis, were ethanol causes intestinal barrier disruption and permits the translocation of microbial products such as lipopolysaccharide (LPS) into the portal circulation (26, 41). This may lead to subsequent activation of hepatic Kupffer cells, exacerbating liver inflammation seen in severe malaria cases (32, 42). The combined effects of increased parasitaemia, immune dysregulation, and liver injury culminate in a more severe disease phenotype, with higher risks of complications such as cerebral malaria, severe anaemia, and multi-organ failure. The purpose of this work is to elucidate the mechanisms via which acute alcohol administration aggravates the pathogenesis of malaria in *Plasmodium berghei*-infected mice by investigating the effects of alcohol on parasitaemia, haematological parameters, liver enzymes, and cytokine levels.

## 2. Materials and Methods

### 2.1. Animals and ethical considerations

Young male Swiss mice (8-10 weeks) with average weight (20-25g) were used for this experiment. They were bred in the Animal house of the Faculty of Science, University of Buea (Cameroon) where they were maintained under ambient temperature (25-27 ^o^C) on a 12-hour light and dark cycle. The animals were fed with conventional commercial diet and tap water *ad libitum*. All experimental procedures were approved by the University of Buea Institutional Animal Care and Use Committee (UB-IACUC) in accordance with the Guidelines for the Care and Use of Laboratory Animals. The approval number of the Ethics Committee was UB-IACUC N°8/2021. Humane endpoints (such as weight loss, behavioural distress, and temperature changes) were applied throughout the experiment and all efforts were made to reduce pain, distress and the suffering of the animals.

### 2.2. Experimental design and animal grouping

In total, 32 male mice were randomly divided into 4 groups: normal control group, alcohol control group, *P. berghei* group and *P. berghei* plus alcohol group. Mice in the *P. berghei*, and *P. berghei* plus alcohol groups were infected with *P. berghei* (10^7^) pRBCs by intraperitoneal injection. Forty-eight hours post infection, mice in the *P. berghei* infected plus alcohol fed group and those in the alcohol control group were administered a single binge concentration of 20% ethanol orally. Control mice received distilled water 10mg/kg body weight. From seventy-two hours post infection, parasitaemia levels were evaluated daily for 3 days. After the evaluation of parasitaemia, animals that survived were subjected to anaesthesia and sacrificed. Blood was collected in tubes with and without anticoagulant (EDTA) by retro-orbital puncture for haematological and biochemical analysis respectively. For biochemical analysis, the blood collected was centrifuged at 3000rpm for 10 min. Serum was separated and stored at -20ºC until used (Figure 1).

**Figure 1.**
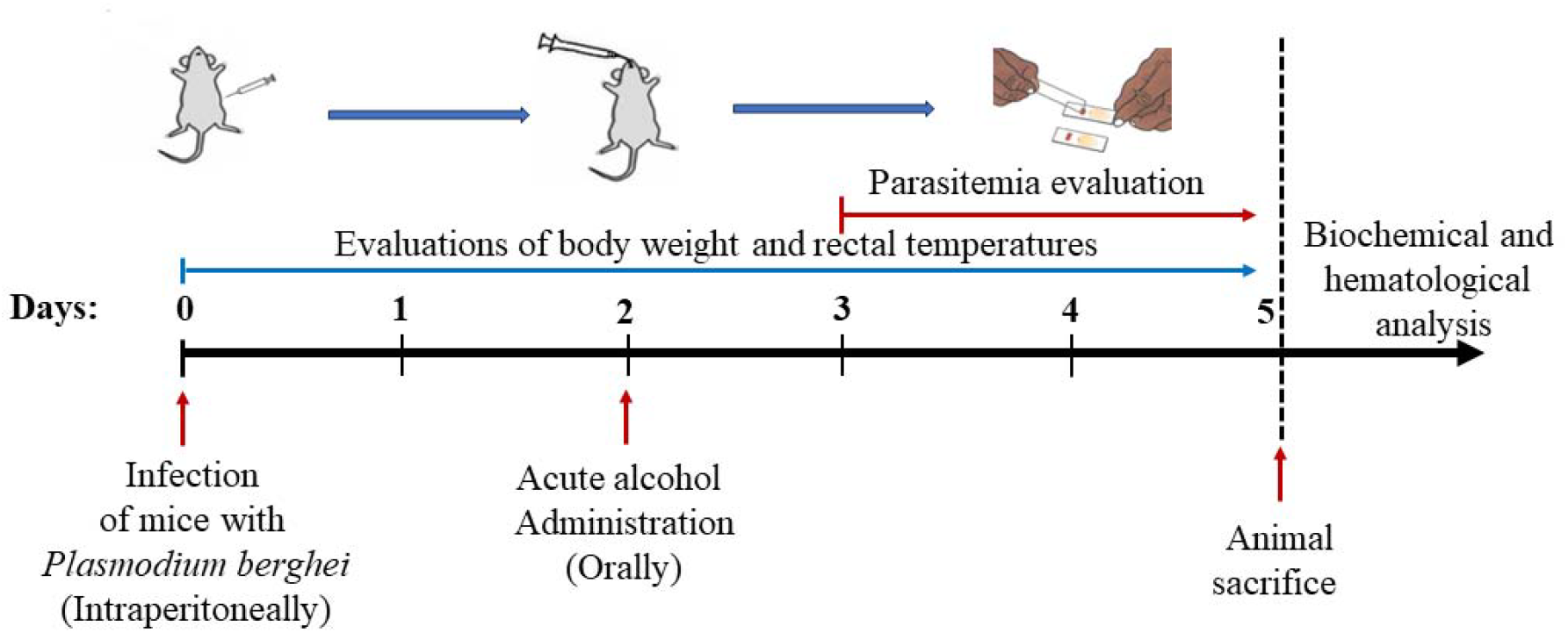
Experimental design

### 2.3. Parasite infection

*Plasmodium berghei* ANKA strain (chloroquine sensitive) was obtained from The Malaria Research and Reference Reagent Resource Centre (MR4, MRA-865, Manassas, Virginia) and stored at -80°C until used. *Plasmodium berghei* ANKA parasite was introduced into a donor mouse by intraperitoneal (i.p) injection of 0.2ml blood containing parasites stored in Alservers buffer solution, and the parasite was allowed to proliferate and establish. The donor mouse was subjected under general anesthesia by inhalation of diethyl ether and blood was collected through cardiac puncture, which was then diluted with sterile normal saline (0.9% NaCl) to obtain 2 x 10^7^ parasitized red blood cells (PRBC). After dilution, blood containing infected red blood cells was obtained. A 0.2 mL blood suspension was injected intraperitoneally into each mouse to cause the infection, which was intended to evolve into a gradually increasing infection. Control mice received an equivalent volume of sterile normal saline.

### 2.4. Chemicals

Reagents used for the determination of AST, ALT and hematological parameters were obtained from Sigma-Aldrich, USA. ELISA kits for cytokine quantification (Quantikine®) were purchased from are from R&D Systems, Minneapolis, USA. Giemsa stain and methanol used for parasitemia determination were also procured from Sigma-Aldrich, USA. Ethanol (95 %) was obtained from a local pharmacy in Buea, Cameroon.

### 2.5. Measurement of body weight and anal temperature

Throughout the study, mice’s body weight was measured using a top pan balance and the values recorded to the nearest 0.1 g. The rectal temperature of mice was measured using a digital rectal thermometer, with the probe carefully inserted through the anal sphincter into the colon of a gently restrained mouse.

### 2.6. Parasitaemia measurement

One drop of blood was collected from the tail of mice via venesection onto the edge of a clean microscope slide (single, 76 × 26 mm thickness). A thin film was prepared by drawing blood evenly across the slide using a second slide an allowed to dry at room temperature before fixing with absolute methanol. The slides were stained with Giemsa stain and viewed using light microscopy technique with oil immersion (1000× magnification). Parasitaemia was evaluated by counting the parasitized red blood cells. Five fields of approximately 200 cells each were counted and the parasitaemia was calculated as the percentage of parasitized red blood cells with respect to the total red blood cells.

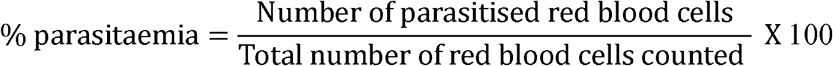

### 2.7. Haematological analysis

Haematological analysis was performed using an automated haematological analyser (Sysmex KX-21N). The haematological parameters evaluated include: red blood cell (RBC) count, leukocyte (WBC) count, haemoglobin (Hb), haematocrit (HCt), package corpuscular volume (PCV), mean corpuscular volume (MCV), mean corpuscular haemoglobin (MCH), mean corpuscular haemoglobin concentration (MCHC), platelet count, lymphocyte, monocyte, neutrophil, basophil and eosinophil counts.

### 2.8. Biochemical analysis

Liver enzymes were evaluated using Sigma diagnostic kits (USA), as described on the manufacturer’s instructions. Aspartate amino transferase (AST) and alanine amino transferase (ALT) activities were assessed in the serum of mice. More so, the concentration of pro-inflammatory cytokines TNF-α, IFN-γ, IL-1β and IL-6 were measured in the serum of mice. All the cytokines were measured by means of ELISA in accordance to Quantikine ELISA kit (R&D Systems, USA) according to the manufacturer’s instructions.

### 2.9. Statistics

Data was analysed and visualized using Graph Pad Prism software version 10 (San Diego, California, USA). Data normality was tested using the Shapiro-Wilk test. Parasitaemia levels in the *P. berghei* and *P. berghei* + Alc. groups were analysed using the RM-ANOVA (Repeated measure ANOVA), followed by Šídák’s multiple comparisons test. Body temperature, body weight, different haematological parameters, liver enzymes (AST and ALT) and pro-inflammatory cytokines were analysed with One-way ANOVA (ordinary) followed by Tukey multiple comparisons test. Results from One-way ANOVA were reported as F-value and p-value. Tukey’s multiple comparisons test results are presented as adjusted p-values. Data were presented as Mean ± Standard error of mean. Mean values with adjusted p values of p < 0.05 were considered significant.

## 3. Results

### 3.1. Parasitaemia development

Parasitaemia levels were evaluated daily for 3 days (from day 3 to day 5) and presented as the percentage of parasitized red blood cells (Figure 2). A non-significant mean effect [F (1, 15) = 2.618, P = 0.127] of time was revealed by a Repeated measure ANOVA, indicating an increase in parasitaemia over time in both *P. berghei* and *P. berghei* + Alc groups. A significant group effect was also revealed [F (2, 15) = 4.526, P=0.029] with higher parasitemia in the alcohol treated parasitized group. A non-significant time × group interaction [F (2, 15) = 0.5505, P=0.588] suggesting that the increase in parasitemia was less pronounced in alcohol-treated parasitized group. Post hoc tests showed that parasitaemia was significantly higher in the alcohol-treated group on days 5 (p = 0.025).

**Figure 2.**
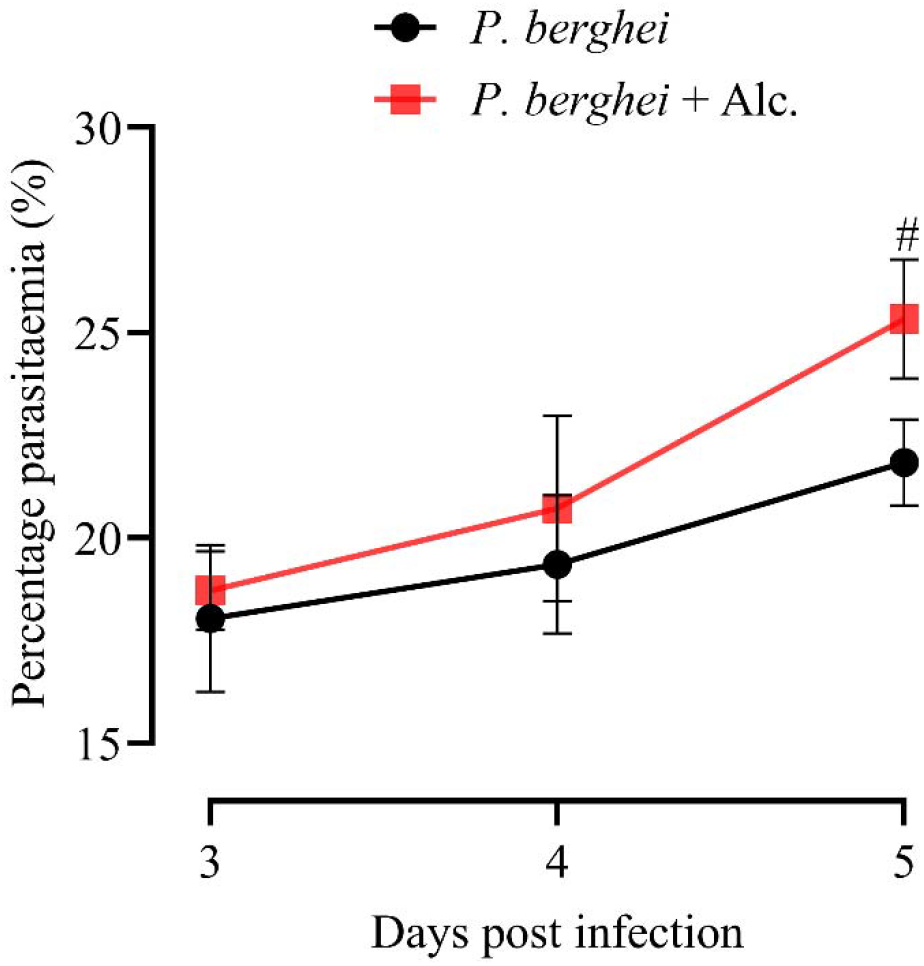
Parasitaemia levels measured in the *P. berghei* and *P. berghei* + Alc. groups. Each line represents the variation of percentage parasitaemia between day 3 and day 5 post infection. Data was expressed as the mean ± S.E.M, n = 8. Data were analysed using RM-ANOVA, followed by Tukey multiple comparisons test; ^#^p < 0.05 versus *P. berghei* group.

### 3.2. Body weight and rectal temperatures

From the results obtained, animals in the control (DW+DW) and the Alcohol control groups progressively gained weight throughout the 5 days of the experiment, whereas, contrarily, a progressive decrease in body weight was observed in *Plasmodium berghei* infected group. This was further worsened by acute alcohol administration to the *Plasmodium berghei* infected mice (*P. berghei* + Alcohol). On day 5, the average body weight was seen to significantly(p<0.05) decrease from 21.51 ± 2.38 g in the *Plasmodium berghei* infected mice to 20.51 ± 1.30 g in the *Plasmodium berghei* infected and alcohol fed mice (Table 1).

**Table 1:** Body weight.

| Day | DW+DW | DW+Alcohol | *P. berghei* | *P. berghei*+Alcohol |
| --- | --- | --- | --- | --- |
| 0 | 24.30 ± 1.55 | 22.07 ± 1.79 | 25.67 ± 2.77 | 24.57 ± 1.16 |
| 1 | 24.90 ± 1.45 | 22.33 ± 1.91 | 25.90 ± 2.70 | 24.03 ± 1.40 |
| 2 | 25.47 ± 1.46 | 23.27 ± 1.75 | 24.80 ± 2.69 | 23.80 ± 1.06 |
| 3 | 25.80 ± 1.45 | 23.8 ± 2.17 | 23.30 ± 2.54 | 23.44 ± 1.36 |
| 4 | 26.00 ± 1.46 | 24.07 ± 1.88 | 22.01 ± 2.19 | 21.07 ± 1.42 * |
| 5 | 26.30 ± 1.65 | 24.68 ± 1.87 | 21.51 ± 2.38 | 20.51 ± 1.30 * |

Data were presented as mean ± S.E.M, n =8. Data were analysed with the ordinary One-Way ANOVA followed by Tukey multiple comparisons test. \* $p < 0.05$ , \*\* $p < 0.01$ , versus body weight of Day 0 in each independent group.

Body temperature changes were observed during the experiment are presented in Figure 3 below. Repeated measure ANOVA shows that there was a significant mean effect [F (2.466, 86.30) = 437.3, P<0.001] of time (in days), indicating a decrease in temperature in the *P. berghei* and *P. berghei* + Alc groups over time. A significant group effect was also observed after repeated measure ANOVA [F (4, 35) = 138.5, P<0.001], with the lowest temperatures observed in the *P. berghei* + Alc groups. There was also a significant time × group interaction [F (12, 105) = 43.98, P<0.001]. Tukey’s multiple comparisons test revealed a significant difference in the *P. berghei* (p < 0.001) and *P. berghei* + Alc (p < 0.001) groups with respect to the control group (DW+DW) (Figure 3).

**Figure 3.**
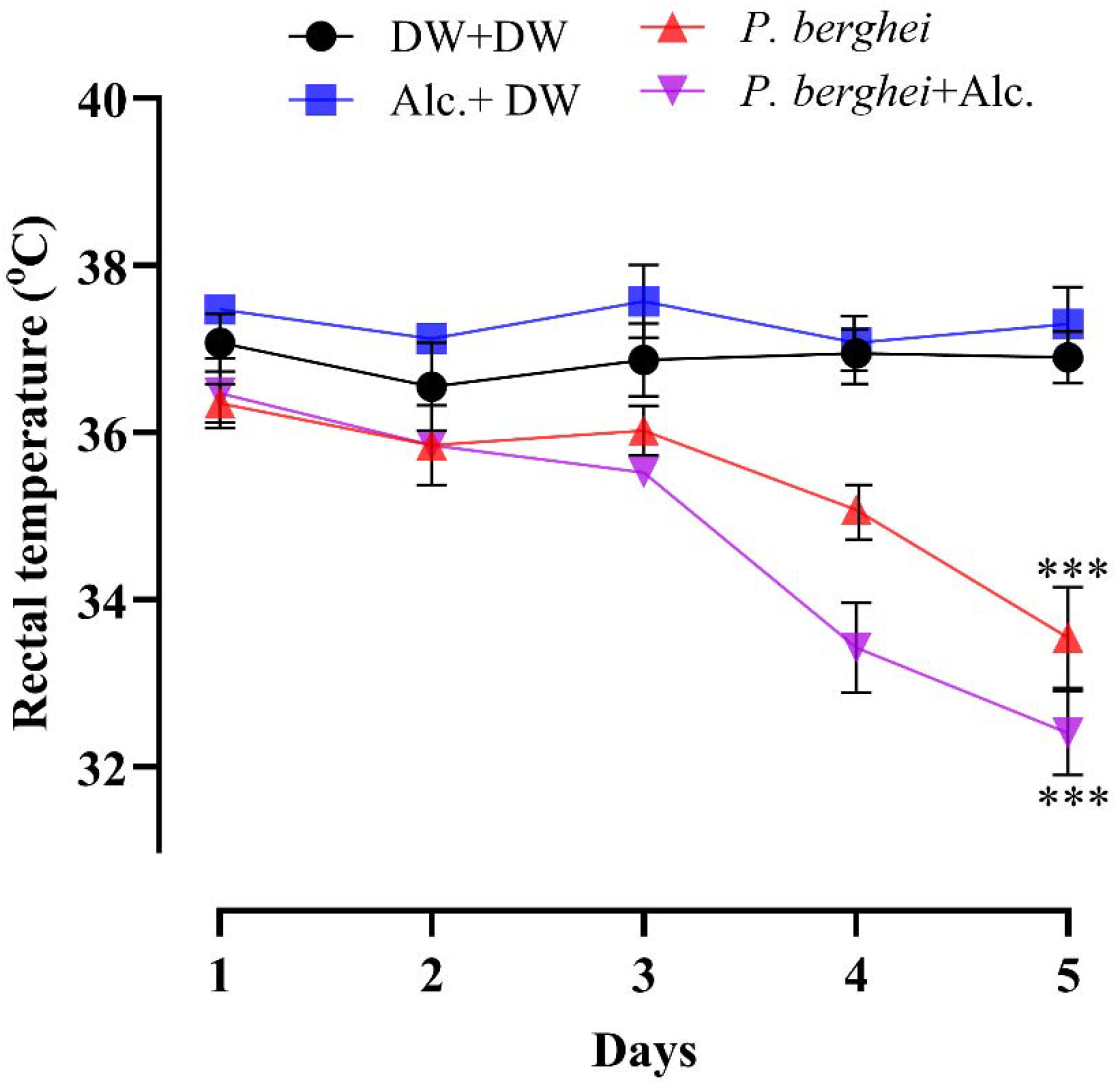
Rectal temperatures of mice in the parasitized and control groups. Each line represents the body temperature variation of each group. Data were presented as mean ± S.E.M, n =8. Data were analysed with the RM-ANOVA followed by Tukey multiple comparisons test. ^***^p < 0.001 versus control group (DW+DW).

### 3.3. Haematological parameters

Table 2 shows the effect of acute alcohol consumption on the different haematological parameters of *Plasmodium berghei* infected mice. As indicated in the results, there was a significant decrease in the level of RBCs [F (3, 20) = 554.4, P<0.001]. The number of RBCs decreased from 6.48 ± 0.09 x106/μL in the control (DW+DW) group of mice to 3.57 ± 0.11 x106/μL (P<0.001) in the *Plasmodium berghei* infected mice and to 2.21 ± 0.13 x106/μL in the *Plasmodium berghei* infected plus alcohol fed group of mice. One-Way analysis of variance revealed significant (P<0.001) reductions in the levels of haemoglobin, haematocrit, MCV, MCH, MCHC, platelets, WBC, neutrophils, eosinophils, lymphocytes, and monocytes in the *Plasmodium berghei* infected group of mice as compared to the control (DW+DW) mice. However, acute alcohol feeding was proven to aggravate the negative effects of *Plasmodium berghei* infection on haematological parameters. As a result of acute alcohol feeding, the levels of RBC reduced from 3.57 ± 0.11 x106 µL^-1^ to 2.21 ± 0.13 x106 µL^-1^, Haemoglobin reduced from 7.53 ± 0.26 g/dL to 4.19 ± 0.26 g/dL, Haematocrit from 24.36 ± 0.35 % to 16.56 ± 1.16 %, MCV from 36.96 ± 0.85 fL to 31.03 ± 2.46fL, MCH from 13.94 ± 0.35 pg to 8.17 ± 0.54 pg, MCHC from 25.04 ± 0.36 g/dL to 21.06 ± 0.5 g/dL, Platelets from 327.7 ± 3.28×103 µL^-1^ to 316.33 ± 3.55 ×103 µL^-1^, WBC from 12.11 ± 0.4 ×103 µL^-1^ to 8.47 ± 0.78 ×103 µL^-1^, Neutrophils from 16.3 ± 0.34% to 12.57 ± 0.67%, Eosinophils from 1.36 ± 0.04 % to 1.33 ± 0.06 %, Lymphocytes from 42.79 ± 3.23 % to 37.78 ± 2.66 % and Monocytes from 2.51 ± 0.12 % to 1.84 ± 0.34 %. However, these reductions were not significant.

**Table 2:**
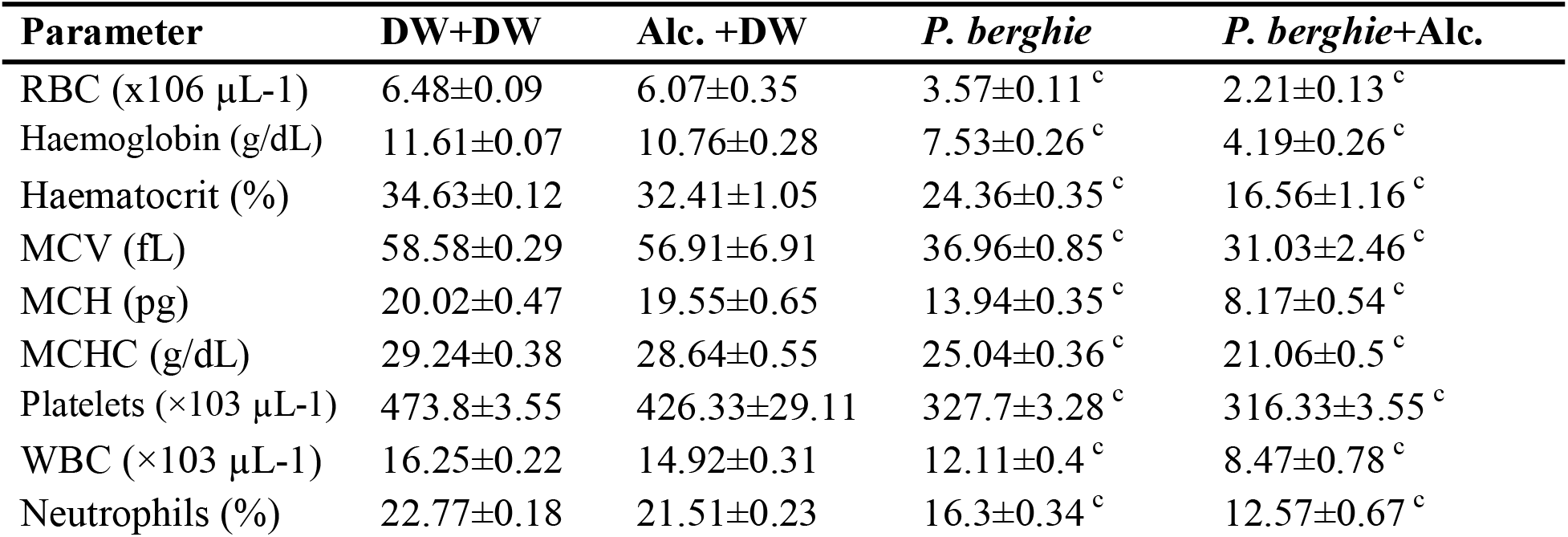
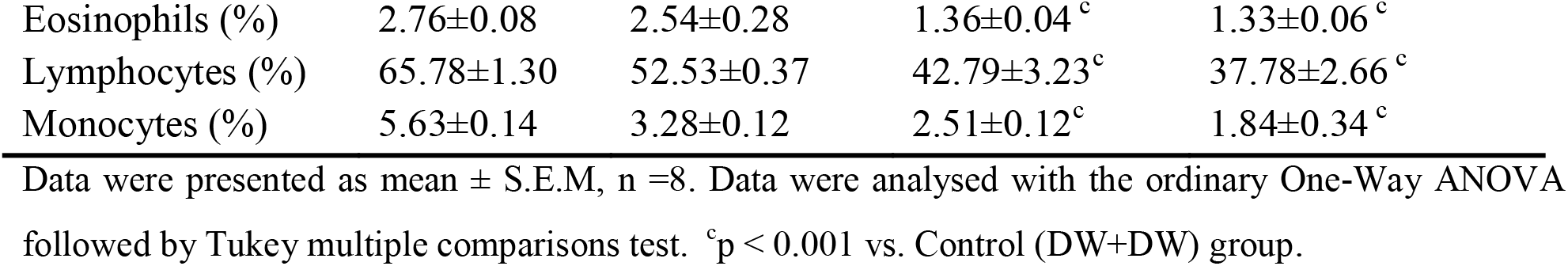
Haematological parameters.

### 3.4. Liver enzymes

After evaluations of liver enzymes, it was shown that *Plasmodium berghei* infection provoked a significant (p<0.001) increase in the activities of aspartate amino transaminase (AST) and alanine amino transaminase (ALT). However, acute alcohol feeding contributed to more activity of AST [F (3, 28) = 36.66, P<0.001] and ALT [F (3, 28) = 29.99, P<0.001] in the serum of mice.

The activities of AST and ALT significantly (p<0.05) increased from 33.39±0.95 U/L and 33.23±1.37 U/L respectively in the *Plasmodium berghei* infected group to 38.02±0.91 U/L and 39.35±1.21 U/L respectively in the *Plasmodium berghei* infected and alcohol fed group (Figure 4).

**Figure 4.**
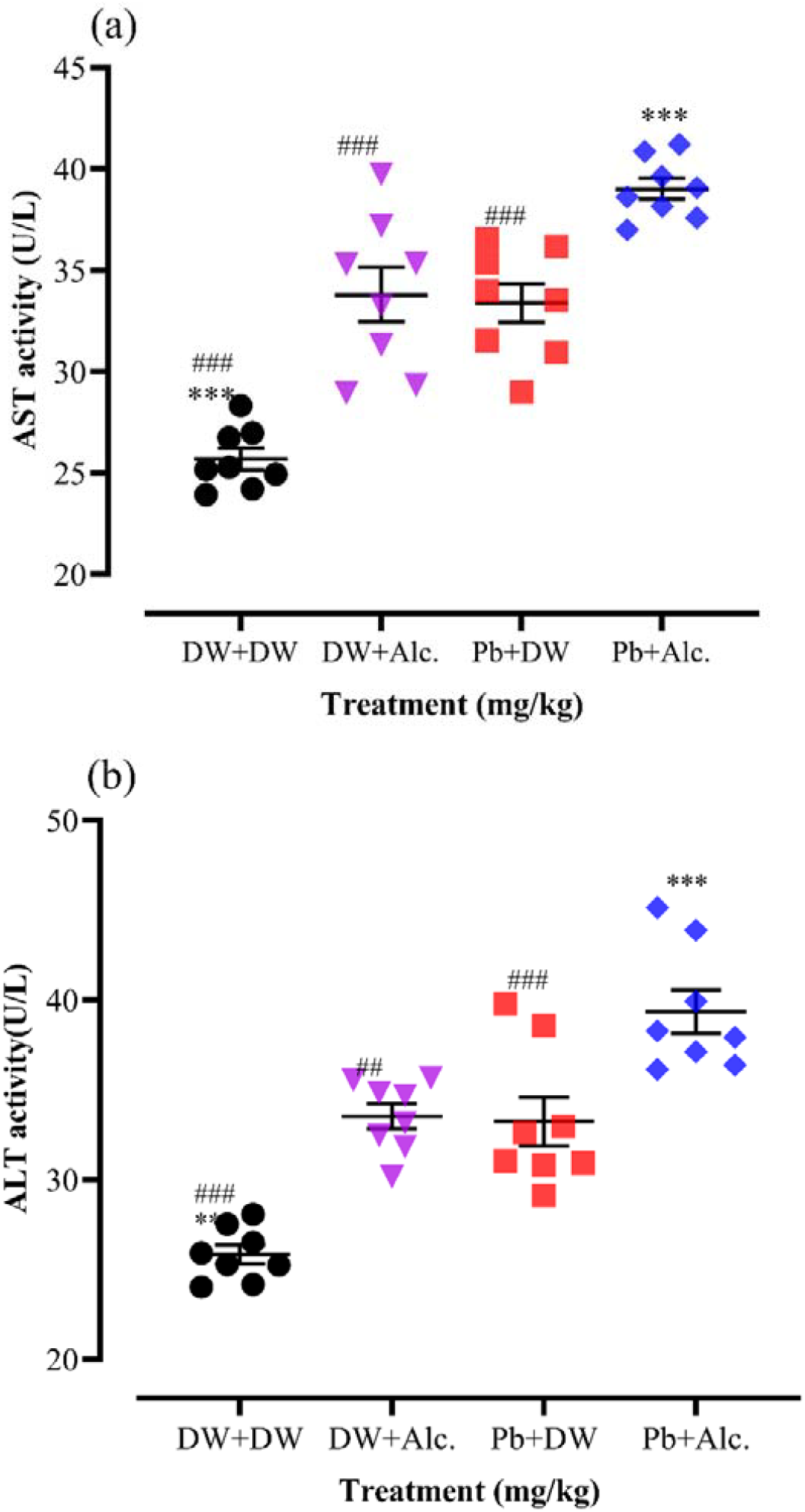
Liver enzymes. Data was presented as individual values in a scatter plot, n = 8. Data were analysed with the ordinary One-Way ANOVA followed by Tukey multiple comparisons test. ***p < 0.001, versus *P. berghei* group, ^###^p < 0.05 versus *P. berghei* plus acute alcohol.

### 3.5. Proinflammatory cytokines

After the evaluation of the concentration of proinflammatory cytokines in the serum of mice, it was observed that *Plasmodium berghei* infection caused a significant (p < 0.001) increase in the levels of TNF-α [F (3, 20) = 170.8, P<0.001], INF-γ [F (3, 20) = 2070, P<0.001], IL-1β [F (3, 20) = 226.1, P<0.001] and IL-6 [F (3, 20) = 82.89, P<0.001] as compared to the control (DW+DW) group of mice. There was a further significant elevation on the concentrations of TNF-α, INF-γ, IL-1β, and IL-6 caused in *P. berghei* infected plus acute alcohol administered group of mice as compared the *P. berghei* only infected group (Figure 5).

**Figure 5.**
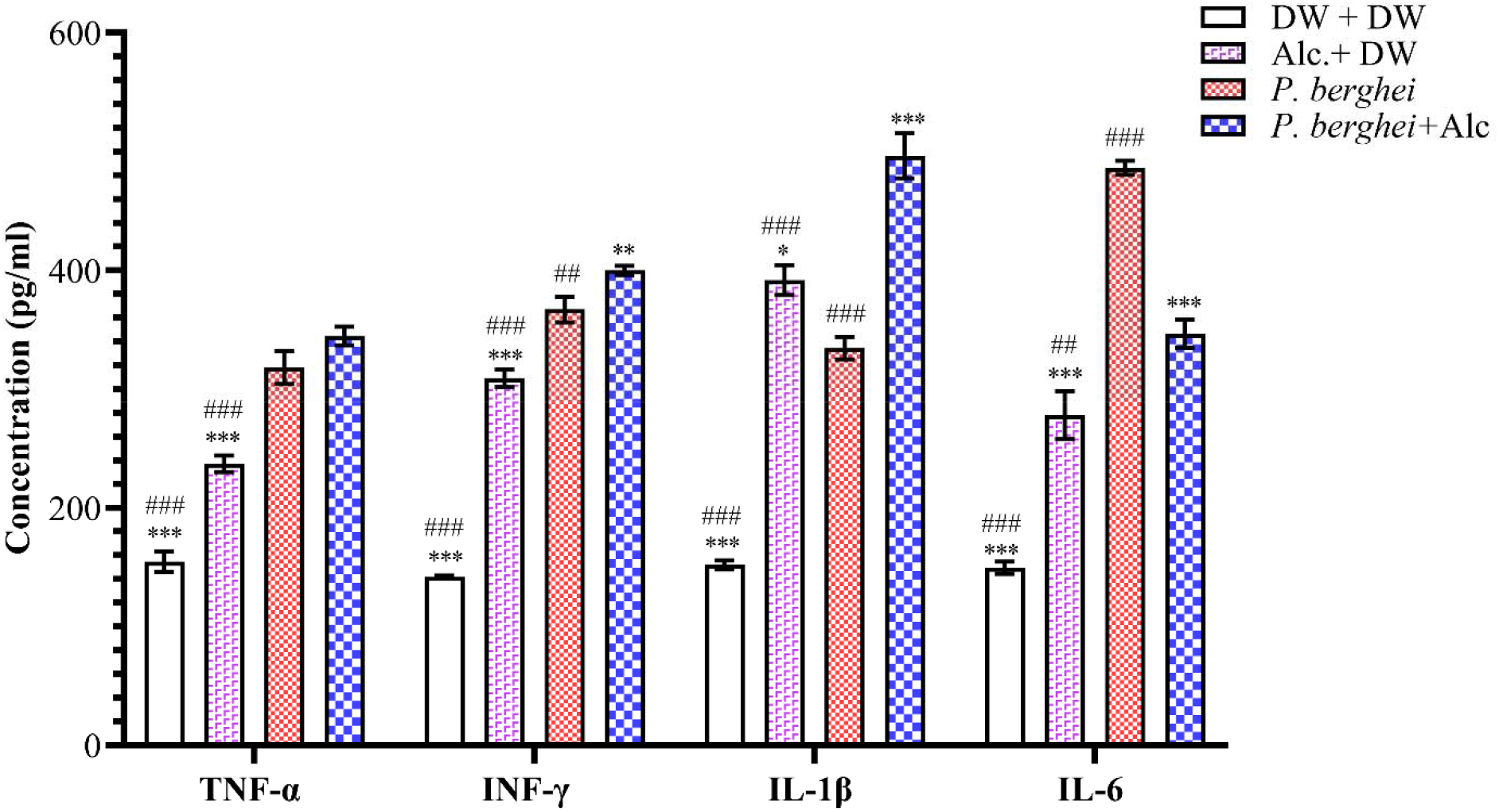
Proinflammatory cytokines. Each bar represents the mean ± S.E.M, n = 8. Data were analysed with the ordinary One-Way ANOVA followed by Tukey multiple comparisons test. ^*^p < 0.05, ^**^p < 0.01, ^***^p < 0.001 *versus P. berghei* group; ^#^p < 0.05, ^##^p < 0.01, ^###^p < 0.001 *versus P. berghei* plus acute alcohol.

## 4. Discussion

The present study sheds light on the crucial insights into the effects of acute alcohol administration on malaria pathogenesis in *Plasmodium berghei*-infected mice. The findings obtained were interesting, and they indicated that acute alcohol administration can exacerbate parasitaemia, haematological parameters, liver enzymes, and cytokine profiles in *Plasmodium berghei*-infected mice. The results obtained confirms our null hypothesis, that ethanol metabolism contributes to intensifying malaria through multiple mechanisms, including metabolic, hepatic and immunological pathways.

A key aspect of this study is that acute alcohol exposure triggers CYP2E1-dependent oxidative stress in the liver, which disrupts key redox-sensitive innate immune pathways which is necessary for early control of *Plasmodium* infection (43, 44). Excessive production of ROS results to impaired hepatocellular homeostasis and attenuates interferon signalling and interferon-stimulated gene expression, weakening restriction of liver stage parasite development in the intracellular level(45, 46).

Concurrently, NF-κB activation in hepatocytes and Kupffer cells is suppressed by alcohol, reducing production of pro-inflammatory cytokines such as TNF-α, IL-6, and INF-gamma that are necessary for effective immune cell recruitment, macrophage activation and early-stage parasite containment(32). Alcohol also induces mitochondrial dysfunction which further disturbs NLRP3 inflammasome signalling, limiting IL-1β and IL-18 maturation and impairs IFN-γ production by NK and T cells(47, 48). Also, direct suppression of Kupffer cell phagocytic and antigen-presenting activities, impairs the clearance of infected erythrocytes and circulating merozoites during the blood-stage infection(49).

Results obtained from this study indicated a significant decrease in body temperature and body weight in the *Plasmodium berghei*-infected groups, including the group exacerbated by acute alcohol administration. The decrease in body temperature can be explained by a combination of host and pathogen-driven mechanisms. During malaria, the hypothermic response is triggered as part of the host’s immune defense strategy (50, 51). This phenomenon is believed to retard the replication of parasites, which have an optimal growth temperature very close to the host’s normal temperature (52). However, this response can be dysregulated under certain conditions, such as co-exposure to alcohol. Alcohol consumption interferes with the hypothalamic-pituitary-adrenal (HPA) axis, impairing the central regulation of body temperature by modifying neurotransmitter release, such as gamma-aminobutyric acid (GABA) and glutamate, which are important in the thermoregulatory pathway (53, 54). Furthermore, alcohol has an impact on prostaglandin synthesis and action, which are important mediators of fever and thermoregulation (55). Prostaglandins, particularly prostaglandin E2 (PGE2), act on the hypothalamus to elevate the body’s set point temperature (56). Alcohol impairs the synthesis of prostaglandins via interfering with the cyclooxygenase enzymes (COX-1 and COX-2) (57), which might exacerbate hypothermia or reduce the fever response.

Similarly, the observed depletion in body weight across the infected groups can be attributed to several factors associated with malaria pathogenesis, including increased metabolic demands due to fever and immune response and anorexia (58). The alcohol-treated malaria group experienced a more severe reduction in body weight, indicating that alcohol consumption further compounds metabolic stress. This could be linked to alcohol’s effect on hepatic function, as metabolism of alcohol in the liver demands a lot of energy, which depletes glycogen stores and puts the body in a catabolic condition.

The present study revealed some haematological alterations, including anaemia, leukopenia, and thrombocytopenia, alongside significant depletion in haematological parameters such as haemoglobin, haematocrit, mean corpuscular volume (MCV), mean corpuscular haemoglobin (MCH), mean corpuscular haemoglobin concentration (MCHC), and differential counts of white blood cells (WBCs), including neutrophils, eosinophils, lymphocytes, and monocytes.

The suppression of bone marrow erythropoiesis by alcohol combined with heightened parasitaemia, which is a condition in which Plasmodium parasites invade and rupture erythrocytes during their life cycle are believed to be the main cause of the observed anaemia, which is characterised by decrease red blood cell count, haemoglobin and haematocrit (38, 59). Changes in MCV, MCH, and MCHC suggest macrocytic anemia caused by vitamin deficiency induced by alcohol (60). Thrombocytopenia can be associated to immune-mediated damage, platelet sequestration, and reduced platelet synthesis which are all aggravated by pro-inflammatory cytokines (61). Leukopenia, which is characterized by significant reduction in neutrophils, lymphocytes, and monocytes, indicates a compromised immune function (62).

Significant rise of liver enzymes, AST and ALT, in alcohol-exacerbated malaria group highlights the hepatotoxic effects of alcohol in the context of malaria. The liver is the key organ for alcohol metabolism and it plays a central role in immune response to malaria, rendering it particularly vulnerable. Acetaldehyde, formed during alcohol metabolism is a highly reactive compound that can cause lipid peroxidation and oxidative stress in hepatocytes(63). This, combined with the inflammatory cytokine storm triggered by malaria infection, could lead to severe hepatocellular damage, evidenced by heightened liver enzymes (ALT and AST). Moreover, although the metabolic shift towards increased glucose synthesis is advantageous for the parasites, it adds extra stress on the liver and exacerbates the damage (64).

Pro-inflammatory cytokines, including TNF-α, IL-6, IL-1β, and IFN-γ, were significantly upregulated in the alcohol-treated malaria group, according to cytokine analysis, which led to a considerable immunological dysregulation. TNF-α and IL-6 contribute significantly to parasite clearance by enhancing phagocytic activity and the generation of inflammatory mediators (16). However, excessive levels of TNF-α and IL-6 are associated to severe pathology, such as cerebral malaria, as a result of tissue damage and disruption of blood-brin barrier (65). Similarly, IL-1β increases the body’s immunological response by triggering fever and immune cell recruitment; however, excessive production of it can result in systemic inflammation and multi-organ damage (65). IFN-γ enhances macrophages’ capacity to phagocytize infected erythrocytes and promotes the production of reactive oxygen and nitrogen species that are toxic to parasites (66). However, chronically elevated levels, especially in conjunction with other pro-inflammatory cytokines, can result in excessive inflammation and tissue damage.

Even though the present study focused on acute alcohol exposure, our findings are consistent with already established mechanisms showing that ethanol metabolism via production of acetaldehyde and activation of NF-κB signaling, induces a biphasic immune response characterized by elevations of pro-inflammatory cytokines such as TNF-α, IL-6, and IL-1β(67), following an early suppression of suppression of innate cytokine. This transient cytokine imbalance disrupts hepatic immune surveillance during the pre-erythrocytic phase of the infection, impairing Kupffer-mediated parasite clearance(68). On the other hand, chronic alcohol exposure is known to produce a sustained immunosuppression via distinct mechanisms(19), however, these effects were not examined in the present study. Thus, our data support a model of acute alcohol potentiating early malaria pathogenesis via oxidative stress and immune dysregulation.

Beyond the directly induced hepatic and immunological effects, acute alcohol exposure can disturb the host’s nutrition/metabolic state, stress/neuroendocrine, and gut–immune axes that have already been shown to modulate malaria pathogenesis. Acute alcohol intake can impair nutrient absorption, impairing glucose metabolism(69), which directly influences *Plasmodium* replication and host resistance. Also, the activation of hypothalamic–pituitary–adrenal axis by acute alcohol exposure induces stress-induced immunomodulation, supressing cytokine production and innate immune responses during early infection(70). Additionally, acute alcohol exposure alters gut microbiota, increasing permeability, and promoting lipopolysaccharide (LPS) translocation into systemic circulation(71). The presence of LPS in circulation can further modulate systemic inflammatory signalling and cytokine levels during blood-stage malaria(72), contributing the impact of acute alcohol on the host-parasite interactions.

The present study had some limitations. Parasitaemia was measured for just 3 days, restricting evaluation of long-term dynamics and survival outcomes, while the use of acute doses limits extrapolation of chronic alcohol effect, which is highly relevant in malaria endemic settings, where alcohol consumption is common. Also, the non-evaluation of IL-10 prevented calculation of IFN-γ/IL-10 ratio to depict immune balance. Additionally, the lack of liver histopathology prevents a firm conclusion on the combined acute alcohol exposure and malaria associated hepatic injury.

However, future studies should incorporate extended periods to monitor parasitemia, chronic and/or binge alcohol models and cytokine profiling to include anti-inflammatory cytokine (such as IL-10).

In conclusion, our findings suggest that acute alcohol consumption may increase malaria susceptibility and severity by interfering with hepatic immune defences and promoting parasite proliferation. These finding underscores the need to consider alcohol intake recent history in malaria treatment and risk assessment, highlighting the value of preventive counselling and supportive strategies that preserve liver functions and immune balance in alcohol consuming populations.

